# The genomic landscape of divergence across the speciation continuum in island-colonising silvereyes (*Zosterops lateralis*)

**DOI:** 10.1101/2020.02.18.953893

**Authors:** Ashley T. Sendell-Price, Kristen C. Ruegg, Eric C. Anderson, Claudio S. Quilodrán, Benjamin M. Van Doren, Vinh Le Underwood, Tim Coulson, Sonya. M. Clegg

## Abstract

A goal of the genomic era is to infer the evolutionary dynamics at play during the process of speciation by analysing the genomic landscape of divergence. However, empirical assessments of genomic landscapes under varying evolutionary scenarios are few, limiting the ability to achieve this goal. Here we combine RAD-sequencing and individual-based simulations to evaluate the genomic landscape in the silvereye (*Zosterops lateralis*). Using comparisons matched for divergence timeframe and gene flow context, we document how genomic patterns accumulate along the speciation continuum. In contrast to previous predictions, our results provide limited support for the idea that divergence accumulates around loci under divergent selection or that genomic islands widen with time. While a small number of genomic islands were found in populations diverging with and without gene flow, in few cases were SNPs putatively under selection tightly associated with genomic islands. Furthermore, we modelled the transition from localised to genome-wide levels of divergence using individual-based simulations that considered only neutral processes. Our results challenge the ubiquity of existing verbal models that explain the accumulation of genomic differences across the speciation continuum and instead support the idea that divergence both within and outside of genomic islands is important during the speciation process.

**DATA ACCESSION NUMBERS:** Resequencing data from this study have been submitted to the National Center for Biotechnology Information (NCBI; https://www.ncbi.nlm.nih.gov) under accession number PRJNA489169.

## INTRODUCTION

Darwin described the process of speciation as a continuum in which differentiation between populations accumulates, first giving rise to well-marked ‘varieties’ or ‘races,’ and in some instances culminating in the formation of new species (Darwin 1859), a view that today is widely accepted (Mallet 2008; Nosil 2012; Powell et al. 2013). Numerous evolutionary processes can promote, stall or reverse trajectory along the speciation continuum, including gene flow, selection, recombination, mutation and drift (Ricklefs and Bermingham 2007; Ravinet et al. 2017; Kearns et al. 2018). Understanding how these processes interact at different stages and how this interaction shapes the speciation process remains a central but challenging goal in evolutionary biology (Butlin et al. 2012; Ravinet et al. 2017). Examining divergence at the level of the genome is a potentially powerful way to address this challenge (Nosil and Feder 2012; Seehausen et al. 2014).

During the divergence process, some regions of the genome diverge rapidly and others more slowly (Seehausen et al. 2014). This variation in the tempo of divergence across the genome results in a heterogeneous genomic landscape (Nosil et al. 2009; Andrew and Rieseberg 2013; Feder et al. 2013; Seehausen et al. 2014; Feulner et al. 2015; Ravinet et al. 2017). During the initial stages of the speciation continuum, divergence is expected to be localised to regions of the genome where loci are under strong divergent selection, forming peaks of divergence often referred to as ‘genomic islands of divergence’ (Feder et al. 2013; Seehausen et al. 2014). As populations move along the speciation continuum, genomic islands are predicted to widen as linkage disequilibrium facilitates divergence of neutral and weakly selected loci via divergence hitchhiking (Nosil et al. 2009; Feder and Nosil 2010). The growth of genomic islands of divergence forms the basis of the divergence hitchhiking model of speciation (Via 2012), which until recently was the prevailing mechanism through which genome-wide levels of divergence were thought to be achieved (Smadja et al. 2008; Via and West 2008). However, the conditions under which divergence hitchhiking can generate large regions of differentiation are limited, requiring small effective population sizes, low rates of migration, and strong selection (Feder and Nosil 2010). As such, alternative mechanisms such as the transition from single locus dynamics to multi-locus coupling (Flaxman et al. 2013, 2014; Butlin and Smadja 2017; Schilling et al. 2018) have been proposed to explain the continued shift from localised to genome wide levels of divergence.

Using closely related taxa at different stages of divergence as a proxy for the speciation continuum, a number of studies have provided evidence that the pattern of genomic divergence accumulates in a way consistent with the mechanisms described above (Martin et al. 2013; Feulner et al. 2015; Supple et al. 2015; Vijay et al. 2016). However, the pace at which divergence accumulates could be accelerated or hindered by other evolutionary processes. For example, gene flow could slow the rate at which divergence accumulates by having a homogenising effect at neutrally evolving or weakly selected loci (Nosil et al. 2017). In contrast, in genetic isolation, divergence proceeds via selection and/or drift, unfettered by this homogenising effect. As such, these different modes of divergence are expected to affect the distribution of genetic divergence values in predictable ways, accounting for the stage of the speciation continuum (Feder et al. 2012) (see Figure 1). To date, comparisons of genomic divergence between races of *Heliconius* butterflies distributed in allopatry versus those in parapatry (Martin et al. 2013) provide the strongest empirical basis for the likely role of gene flow in shaping the genomic landscape. However, studies are needed with comparisons matched for divergence timeframe and with rates of gene flow inferred from more than geographic distribution alone.

**Figure 1:**
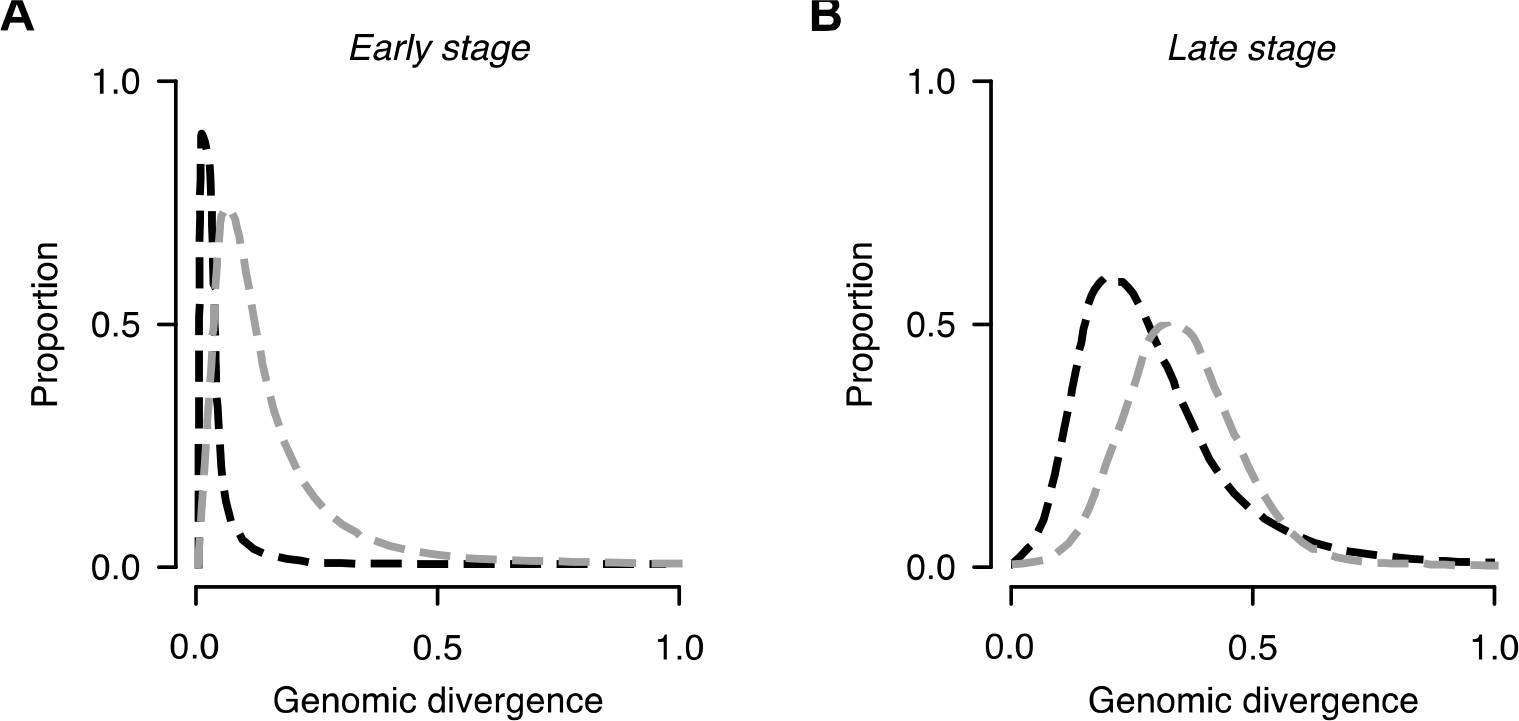
Hypothetical distributions of genetic differentiation expected for divergence with and without gene flow. **(A)** During the early stages of the speciation continuum divergence is expected to be limited to few loci, resulting in a highly skewed distribution of divergence values. This expectation will be most extreme for populations diverging with gene flow (black lines) than for populations diverging in genetic isolation (grey lines) as the homogenising effect of gene flow limits divergence to only loci under strong divergent selection. **(B)** During later stages of the speciation continuum skew is expected to break down as divergence accumulates, but less so in populations diverging with gene flow (black lines) than those diverging without (grey lines) as occasional gene flow may reduce levels of divergence at positions of the genome even as reproductive isolation is approached.

Genomic valleys of similarity – highly conserved regions where differentiation falls far below background levels - are also a common feature of the genomic landscape. While these have been identified in a range of species (Hofer et al. 2012; Roesti et al. 2012; Wang et al. 2016; Van Doren et al. 2017), their role in shaping the genomic landscape has been overlooked in comparison to the focus on genomic islands. Genomic valleys may play an important role in maintaining genomic heterogeneity by slowing the approach to genome-wide divergence. Genomic valleys may occur because of alleles favoured in both populations (parallel adaptation) (Nielsen 2005; Roesti et al. 2012), or they may result from incomplete lineage sorting at neutral loci, where diverging populations share alleles for some time, manifesting as regions of below background level divergence (Stölting et al. 2013). Should parallel adaptation be important in their formation, we would expect the position of genomic valleys to closely correspond to the position of loci under parallel selection. Further, as parallel selection leaves a distinct signature of reduced diversity, where genomic islands are the product of parallel selection, between population diversity (*d*_xy_) would be expected to be reduced (Roesti et al. 2014). Where genomic valleys are the product of incomplete lineage sorting at neutral loci, they are expected to be more numerous during the early stage of divergence as shared alleles become less numerous over time. Under these proposed mechanisms, a reasonable expectation of the temporal dynamics of genomic valleys is that they decrease in size over longer divergence timeframes, as genomic valleys are broken down by recombination and loosely linked/neutral loci diverge (Charlesworth 2006). This expectation is yet to be tested empirically.

A more nuanced understanding of how genome-wide divergence develops from the heterogeneous genomic landscape, and the dynamics of genomic islands of divergence and genomic valleys of similarity, requires simultaneous consideration of the stage along the speciation continuum and the gene flow context of divergence for systems that have well characterised population histories. Members of the silvereye subspecies complex (*Zosterops lateralis*) of Australia and southwest Pacific islands offer an exceptional opportunity to explore patterns of divergence across different timescales and gene flow contexts. The species has repeatedly colonised islands in the region since its origin on the Australian mainland (Mees 1969) across timeframes from hundreds of years to hundreds of thousands of years. This provides a spectrum of divergence timeframes that can be used as a proxy for the speciation continuum, capturing incipient phenotypic divergence through to highly divergent subspecies (Clegg, Degnan, Kikkawa et al., 2002). Importantly, silvereye populations have diverged under different gene flow scenarios, with some populations diverging with gene flow and others diverging in genetic isolation. Further, as insular silvereyes show a repeated pattern of phenotypic change towards increased body size and bill size, largely driven by directional natural selection (Mees 1969; Clegg, Degnan, Moritz, et al. 2002; Clegg et al. 2008), genomic islands may form around loci underlying this phenotypic change across comparisons. While we do not know the genetic architecture of these traits, several different studies have found genes of large effect for bill traits (Cornetti et al. 2015; Lamichhaney et al. 2015; Chaves et al. 2016; Bosse et al. 2017; vonHoldt et al. 2018).

Here we use empirical analysis of Restriction-site Associated DNA sequencing (RAD-Seq) data from silvereyes and individual-based simulation models to address the following questions: 1) How does genomic divergence accumulate across the genome over time? 2) How is the accumulation of divergence affected by gene flow? 3) Does the position of genomic islands of divergence and valleys of similarity correspond to the location of loci under divergent selection or parallel/balancing selection, respectively? 4) Do genomic islands widen, and valleys narrow, over longer divergence timeframes? and 5) Are the features of genomic valleys consistent with proposed mechanisms that could contribute to their formation?

## MATERIALS & METHODS

### Sample collection

Blood samples were collected from nine silvereye populations across the south Pacific, capturing a range of divergence levels and gene flow variation within this species, and providing five diverging population pairs for comparison. Population pairs varied in their divergence timeframes (early stage: <150 years, mid stage: 3,000-4,000 years, and late stage: 100,000s years) and mode of divergence (gene flow or no gene flow) (Figure 2). The populations used in this study provide a rare example of where gene flow scenarios and divergence timescales are known *a priori* or where a very strong case can be made to infer the gene flow scenario. This is based on a combination of historical records, contemporary bird movements, and phylogenetic inferences as outlined below:

**Figure 2:**
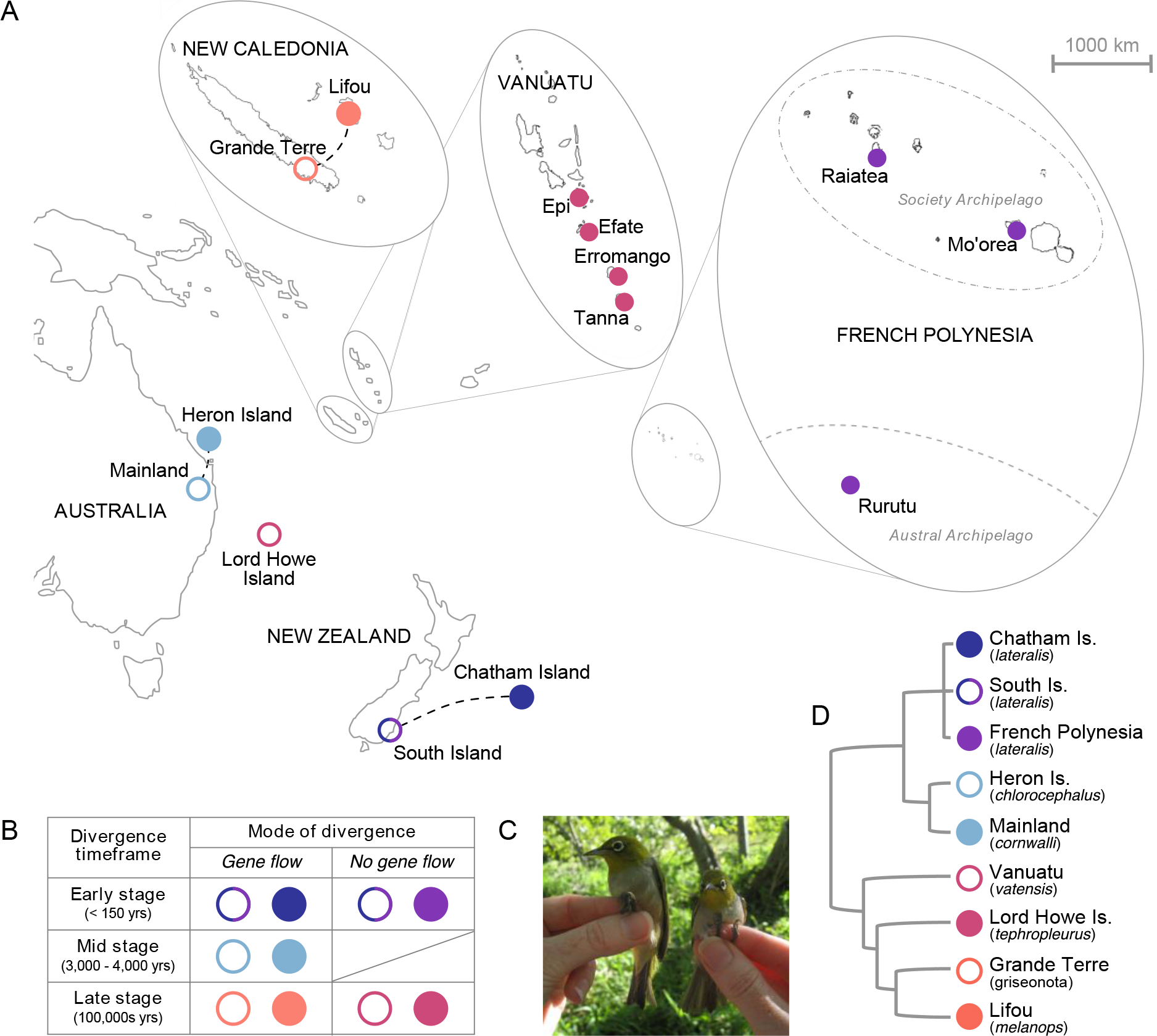
Silvereye populations used in the study. (A) Sampling locations of *Zosterops lateralis* across the south Pacific (dashed lines indicate gene-flow between populations); (B) Population divergence timeframes (known from historical records, or inferred from island ages and genetic divergence dates) and modes of divergence (gene-flow or no gene-flow); (C) *Z. l. chlorocephalus* from Heron Island (left) are up to 40% larger than *Z. l. cornwalli* from the Australian mainland (right). Image by Nick Clark; (D) evolutionary relationships between *Z. lateralis* subspecies and populations based on mitochondrial DNA (Black 2010) and known colonisation histories (subspecies indicated in brackets).

#### South Island vs. Chatham Island

The Chatham Island silvereye population was colonised from South Island (New Zealand) during the 1850s. This timing is historically documented (Mees 1969; Clegg, Degnan, Kikkawa, et al. 2002) and reflected in population level phylogenies (Black, 2010). In the mid 1800s, South Island silvereyes were undergoing a population expansion which resulted in the colonisation of other islands in the region (notably: North Island of New Zealand and Norfolk Island) (Mees 1969; Clegg, Degnan, Kikkawa, et al. 2002). Given this recent colonisation date and that population expansion was taking place within the region, this comparison is characterised as Early Stage – Gene Flow.

#### South Island vs. French Polynesia

The French Polynesia silvereye population is the product of a documented human-mediated introduction of South Island silvereyes (*Z. l. lateralis*) to Tahiti in 1937 (Monnet et al. 1993; Thibault and Cibois 2017). Given that French Polynesia is located well beyond the natural distribution limit for this species and that the French Polynesian population is known to be the product of a single and recent introduction event, this comparison is characterised as Early Stage – No Gene Flow.

#### Mainland (Australia) vs. Heron Island

The Heron Island silvereye population was established from the mainland silvereye population between 3,000-4,000 years ago. This divergence timeframe is supported by geological records which suggest that Heron Island has been vegetated (and therefore habitable) for a maximum of 4,000 years (Hopley 1982; Degnan and Moritz 1992). Given that Heron Island is located only 80km off the Queensland coast and that the ancestral mainland subspecies (*Z. l. cornwalli*) is regularly observed on the island, this comparison is characterised as Mid Stage – Gene flow.

Grande Terre vs. Lifou: Based on mitochondrial DNA divergence estimates, silvereyes on the islands of Grande Terre (*Z. l. griseonotus*) and Lifou (*Z. l. melanops*) diverged in the past few hundred-thousand years (Black 2010). Given the silvereye’s ability to cross water boundaries and the geographic proximity of these islands (200km apart), this comparison is characterised as Late Stage – Gene Flow.

Vanuatu vs. Lord Howe Island: Based on mtDNA phylogenetic analyses, *Z. l. tephropleurus* on Lord Howe Island and *Z. l. vatensis* from Southern Vanuatu are thought to have diverged in the past few hundred-thousand years (Black 2010). Given the geographic isolation of Lord Howe Island, this comparison is characterised as Late Stage – No Gene Flow.

Sampling locations and collection dates are provided in Table 1. To provide large enough sample sizes for analysis, French Polynesian and Vanuatu populations included samples taken from multiple islands (3 and 4 respectively). This is justified based on population genetic clustering of these island populations (Supplementary Figure S1 and Figure S2). All birds were caught using mist nets or traps and 20–40 μl of blood collected from the brachial wing vein was stored in ∼0.5 ml of lysis buffer (0.01M Tris-HCl; 0.01M NaCl; 0.01M EDTA; 1% n-lauroylsarcosine, pH 8.0) (Seutin et al. 1991).

**Table 1:** Z. lateralis sampling information. No. Sequenced = number of individuals including in RAD-Seq libraries; No. Filtered = number of individuals retained after quality filtering.

| Subspecies | Pop. code | Sampling location | Lat / Long | Collection date | No. Sequenced | No. Filtered |
| --- | --- | --- | --- | --- | --- | --- |
| *chlorocephalus* | HI | Heron Island | -23.44 / 151.91 | 2015 | 20 | 16 |
| *cornwalli* | ML | Mainland Australia | -28.18 / 153.45 | 2013 | 20 | 18 |
| *griseonotus* | GT | Grande Terre | -21.78 / 166.02 | 2014 | 20 | 19 |
| *lateralis* | CI | Chatham Island | -44.58 / 176.32 | 1997 | 22 | 18 |
| *lateralis* | SI | South Island | -45.53 / 170.30 | 1997 | 23 | 21 |
| *lateralis* | FP | Mo'orea | -17.51 / -149.91 | 2005 | 17 | 11 |
| *lateralis* | FP | Raiatea | -16.82 / -151.46 | 2010 | 3 | 2 |
| *lateralis* | FP | Rurutu | -22.48 / -151.34 | 2012 | 3 | 3 |
| *melanops* | LF | Lifou | -20.89 / 167.25 | 2014 | 20 | 19 |
| *tephropleurus* | LH | Lord Howe Island | -31.33 / 159.05 | 1998 | 20 | 19 |
| *vatensis* | VN | Epi | -16.74 / 168.27 | 2004 | 1 | 1 |
| *vatensis* | VN | Efate | -17.66 / 168.43 | 2004 | 5 | 4 |
| *vatensis* | VN | Erromango | -18.82 / 169.16 | 2004 | 12 | 9 |
| *vatensis* | VN | Tanna | -19.53 / 169.27 | 2004 | 6 | 4 |

### RAD-PE sequencing and bioinformatics

Restriction-site associated DNA paired-end (RAD-PE) sequencing was used to assess the genomic landscape of divergence among silvereye populations. This reduced representation sequencing method has the potential to identify many hundreds of thousands of single nucleotide polymorphisms (SNPs) distributed throughout the genome (Davey and Blaxter 2010). Genomic DNA in blood samples was extracted from between 20 and 24 individuals per population (or set of populations in the case of Vanuatu and French Polynesia) (n = 192) using QIAGEN DNeasy blood and tissue extraction kits following manufacturer protocols and RAD-PE libraries prepared following the protocol by Ali et al. (2016). See supplementary methods for a comprehensive description of library preparation. Libraries were sequenced on three Illumina HiSeq4000 lanes (Illumina, San Diego, CA, USA) at the UC Davis Genome Center using paired-end 150-bp sequence reads.

Quality of sequencing reads was checked visually using FASTQC (Andrews 2010). The *process_radtags* script included in the STACKS version 1.4 software pipeline (Catchen et al. 2013) was used to assign sequence reads to individuals. In addition, reads containing uncalled bases and/or bases of low quality were discarded in this step using default quality thresholds (an average Phred score of 10 in sliding windows of 15% of the length of the read). Sequences with possible adapter contamination and/or missing the *Sbf1*-HF restriction site were also discarded. Following this, reads were filtered for PCR duplicates using the STACKS *clone_filter* script. The remaining reads were then mapped to the *Zosterops lateralis melanops* genome assembly version 1 (NCBI Assembly GCA001281735.1, (Cornetti et al. 2015)) with BOWTIE2 version 2.2.6 (Langmead and Salzberg 2012) using end-to-end alignment and default settings (allowing for a maximum of two mismatches in the seed (-n 2)). Genotypes were then called using the *HaplotypeCaller* and *GenotypeGVCFs* tools from the Genome Analysis Toolkit (GATK) nightly build version 2016-12-05-ga159770 (McKenna et al. 2010). To output both variant and non-variant sites, *HaplotypeCaller* was ran using -output_mode ’EMIT_ALL_CONFIDENT_SITES’ and *GenotypeGVCFs* ran using mode ’--includeNonVariantSites’. The resulting output was then filtered to remove indels and only include sites where the minor allele count was ≤ 2; minimum genotype quality = 30; minimum depth = 8; and sites were called in at least 70% of individuals. Following this the outputted VCF file was further filtered to remove individuals missing > 30% of sites.

As the *Z. l. melanops* genome is only assembled to the scaffold level (Cornetti et al. 2015), *Z. l. melanops* scaffolds were mapped to chromosomes of the *Taeniopygia guttata* genome assembly version 3.2.4 (NCBI Assembly GCA_000151805.2) using Satsuma Synteny (Grabherr et al. 2010). Output from Satsuma Synteny was then used to assign scaffolds to chromosomes and determine order, location, and orientation using custom R scripts from Van Doren et al. (2017). Further custom scripts were used to reorder the GATK outputted VCF file accordingly. Because synteny is high in birds (Ellegren 2010), 96.8% of the *Zosterops* scaffolds were assigned to assembled chromosomes and arranged in the presumed correct order and orientation. As inversions and other chromosomal rearrangements are known to occur in birds (Backström et al. 2010), it may be possible that a small percentage of scaffolds were ordered or oriented incorrectly. However, a small number of misplaced scaffolds is not expected to impact our overall interpretation of empirical patterns.

Finally, Wright’s fixation index (*F*ST - a measure of relative divergence) and Nei’s measure of absolute divergence (*d*_xy_) were calculated in non-overlapping windows of 50kb and 500kb for each population comparison, using Python scripts developed by Martin et al. (2013) (https://github.com/simonhmartin/genomics_general).

### Population genomic analyses

We assessed the shift from localised divergence at few loci towards more genome-wide levels of divergence by comparing the third moment (skewness) of *F*ST distributions between population comparisons. In line with expectations outlined in Figure 1, skewness provides a useful metric to describe a population comparisons’ stage of transition from localised (highly skewed distributions) to more genome-wide levels of divergence (less skewed distributions). We tested for significant differences in the skew of empirical distributions using a randomisation test, in which we compared absolute differences in skew for observed distributions to absolute differences calculated for 10,000 randomised distributions.

R scripts, based on those used in Van Doren et al. (2017), were used to identify highly diverged regions (genomic islands of divergence) and regions of low divergence (genomic valleys of similarity) occurring across the genomic landscapes of the diverging populations. First, for each population comparison, a kernel-based smoothing algorithm was applied to *F*ST values calculated across the genome in 50kb non-overlapping windows, and the smoothed line compared to 5,000 smoothed lines obtained after permuting the order of the windows (see Ruegg et al. (2014)). We used a window size of 50kb for genomic island and genomic valley detection because it provided a sufficiently fine resolution across the genome while containing on average 1,093 sites per window. Further, windows containing <10 sites were removed prior to conducting analysis. Genomic islands were identified as any location where the observed smoothed line was greater than the most extreme value from the permutation distribution (see Van Doren et al. (2017)). In contrast, genomic valleys were identified as any region where the observed smoothed line was lower than the lowest value of the permuted distribution. Adjacent outlier windows considered in the same genomic island or genomic valley were merged into larger divergent regions. For each genomic island and genomic valley, we calculated mean *F*ST and *d*_xy_. Wilcoxon signed-rank tests were used to determine if *d*_xy_ within genomic islands and genomic valleys differed significantly from chromosomal background levels. To assess the effect of time and gene flow on the development of patterns of heterogeneity the number and size of identified genomic islands and valleys were compared across timescales and gene flow contexts using Kruskal-Wallis tests.

### Identifying outlier SNPs under selection

To identify genomic regions that may be under selection, we scanned for outlier loci using *PCAdapt*, a principal components-based method of outlier detection with a low rate of false-positive detection (Luu et al. 2017). *PCAdapt* requires selection of K principal components, based on inspection of a scree plot, in which K is the number of PCs with eigenvalues that depart from a straight line. *PCAdapt* then computes a test statistic based on Mahalanobis distance and controls for inflation of test statistics and false discovery rate (FDR). Outlier SNPs were identified using the following settings for all population comparisons: K = 2, MinMAF= 0.2, and FDR = 0.001.

### Candidate gene analysis

To determine if genomic islands of divergence occurred around genes known to be associated with body and beak size differences in birds, we compiled a list of candidate genes based upon a literature review of gene expression and association studies. The resulting candidate gene list was restricted to only those genes where (1) locations within the zebra finch genome are known; and (2) genes were mapped to directly by SNPs sequenced as part of this study or genes were located within the 50kb windows used when summarising divergence statistics. Positions of candidate genes were then compared to the position of identified genomic islands of divergence. The position of candidate genes was also compared to the position of outlier SNPs. Genes were determined to fall within genomic islands, if island bounds (start and end positions) overlapped gene start and end positions (this includes 5’ UTRs and introns). Likewise, outlier SNPs were determined as within candidate genes if they fell within the gene start/end positions.

### Simulations of divergence

We performed individual-based simulations where genomic divergence could only occur via neutral processes and compared model predictions with empirically observed genomic patterns. The simulations consequently provide a neutral model. If such a model can generate patterns observed in nature, then it is not necessary to invoke selection as the underlying driver of observed patterns. In other words, the simulation modelling allowed us to address whether the transition from localised to more genome-wide divergence could be explained by drift alone, and also allowed us to explore the fine scale effects of gene flow and recombination by simulating divergence under various levels of gene flow and recombination (both of which were not possible with empirical comparisons alone).

Each simulation considered two populations diverging with or without gene flow. Each population started with individuals with randomly drawn genotypes across 8,317 biallelic SNPs on chromosome 5. Each genotype was determined by sampling with replacement from the distribution of all observed genotyped birds at each SNP. Individuals were then assigned a sex assuming a 50% sex ratio and assigned to a population to produce two populations, each with an initial population size of 400 individuals (similar to the Heron Island breeding population size (Kikkawa and Wilson 1983)).

Each population was iterated forward on a per-generation time step. At each generation in the simulation, individuals were assigned a number of offspring independent of genotype. Fitness (the number of offspring) was determined by randomly drawing from a Poisson distribution with a mean that varied with population density. We thus defined mean population fitness as the average number of offspring produced. We included density-dependence in this function to avoid exponential growth. The density dependence kept the population within empirically observed bounds for the Heron Island population (100-300 breeding pairs). Male fitness was then scaled such that the total sum of male fitness equalled the total sum of female fitness. Mating pairs were formed by combining male and female parents as a function of their ranked fitness values. This approach assumes monogamous reproduction which is empirically observed (Robertson et al. 2001).

To determine offspring genotype, we first identified crossover points along the parental genomes by drawing random values from an exponential distribution. The crossover points assigned along the chromosomes were compared to the genotype structure of individuals and the position of each SNP. Recombination occurred when the random variable was less than a particular recombination rate. Four different recombination rates were simulated: 2 cM/Mb, 3 cM/Mb, 6 cM/Mb, 10 cM/Mb and 19 cM/Mb (translated to crossovers per base pair in our simulations). This corresponds to the range of values known from the collared flycatcher (*Ficedula albicollis*) genome (Kawakami et al., 2014) (Table S1). Flycatcher recombination rates were used, as unlike the zebra finch genome, the flycatcher is not thought to contain large recombination deserts (Backström et al., 2010). Copies of chromosome 5 were then segregated from the parental genomes with recombination occurring at each crossover. Migration rate (*m*) was set to zero in simulations of no-gene-flow scenarios. In divergence-with-gene-flow simulations, offspring either stayed in the same population or dispersed, with simulations repeated for values of *m* of 0.0001, 0.001 and 0.01. Simulations were run for a maximum of 2,000 generations (assuming a generation time of three years (Clegg et al., 2008). Simulations were run in R as described in Quilodrán et al. (Quilodrán et al., 2019). R code is available at GitHub: https://github.com/eriqande/gids.

## RESULTS

### RAD-PE sequencing and bioinformatics

Overall, RAD-PE sequencing resulted in 1,149,472,877 paired-end reads. Post-quality filtering (removal of indels and only including sites where: the minor allele count was ≤2; minimum genotype quality = 30; minimum depth = 8; and genotypes were called in at least 70% of individuals) reads covered a total of 15,872,783 sites (of which 435,456 were biallelic). Of the 192 samples sequenced, 150 were retained after removing individuals where ≥30% of data was missing. The number of individuals retained per location ranged from 16 to 21 (Table 1).

### Overall levels of divergence

Mean *F*ST was lowest for populations in the earliest stages of divergence (South Island vs. Chatham Island and South Island vs. French Polynesia) and increased over longer divergence timeframes, with populations at later stages of divergence (Lord Howe vs. Vanuatu and Lifou vs. Grande Terre) having the highest mean *F*ST (Figure 3A). Mean *F*ST was also consistently lower for populations diverging with gene flow when compared to non-gene flow scenarios matched for divergence timeframe (i.e. South Island vs. Chatham Island compared to South Island vs. French Polynesia) demonstrating how the homogenising nature of gene flow impedes divergence. Compared to autosomes, chromosome Z exhibited higher levels of divergence for all population comparisons (Figure 3A). This difference was significant for South Island vs. Chatham Island, Heron Island vs. Mainland, and Lord Howe vs. Vanuatu comparisons (Mann-Whitney: all *P* values <0.05).

**Figure 3:**
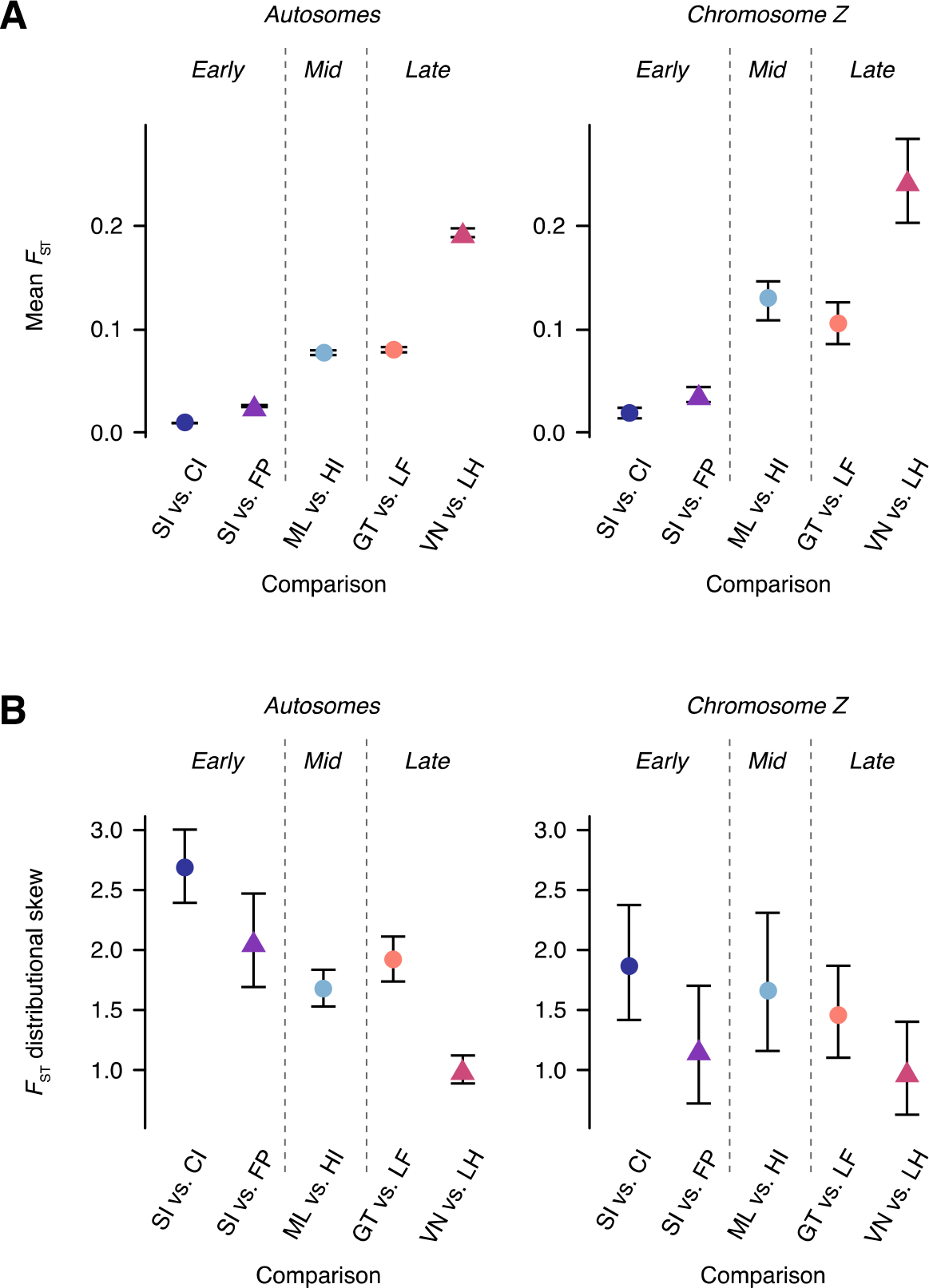
Mean FST and distributional skew of F_ST_ values. **(A)** Mean *F*_ST_ and **(B)** distributional skew of *F*_ST_ values. Calculated for autosomes and chromosome Z separately for each population comparison. Populations diverging with gene flow are indicated by circles and populations diverging in isolation by triangles. 95% confidence intervals obtained via bootstrapping over loci.

### Accumulation of genomic divergence across the genome

In the early stage of the speciation continuum (South Island vs. Chatham Island and South Island vs. French Polynesia comparisons), the distributions of *F*ST values calculated in 50kb windows for autosomes were highly skewed towards large values of *F*ST (skew = 2.69 and 2.07 respectively) (Figure 3B) and characterised by extreme L-shaped distributions with divergence limited to few 50kb windows (Figure 4B). At later stages of the speciation continuum distributional skew decreased with the lowest levels of skew (skew = 1.00 and 1.27) observed for populations in the late stage of divergence (Vanuatu vs. Lord Howe Island and Lifou vs. Grande Terre) and intermediate levels of skew (skew = 1.67) observed for the single mid stage comparison (Mainland vs. Heron Island) (Figure 3B). The accumulation of genome-wide divergence over longer divergence timeframes is clearly visible in Figure 4A. When matched for divergence timeframe, the *F*ST distributions of populations diverging under the influence of gene flow showed higher levels of skew (more localised divergence) than populations diverging in the absence of gene flow. Although not all pairwise differences were significant, recently diverging populations had significantly higher skew for autosomes than late stage diverging populations (Table 2). When treating each autosomal chromosome as an independent unit, more chromosomes showed significantly reduced *F*ST distributional skew than increased skew when diverging under gene flow, during both early and late stage divergence (Table S2).

**Figure 4:**
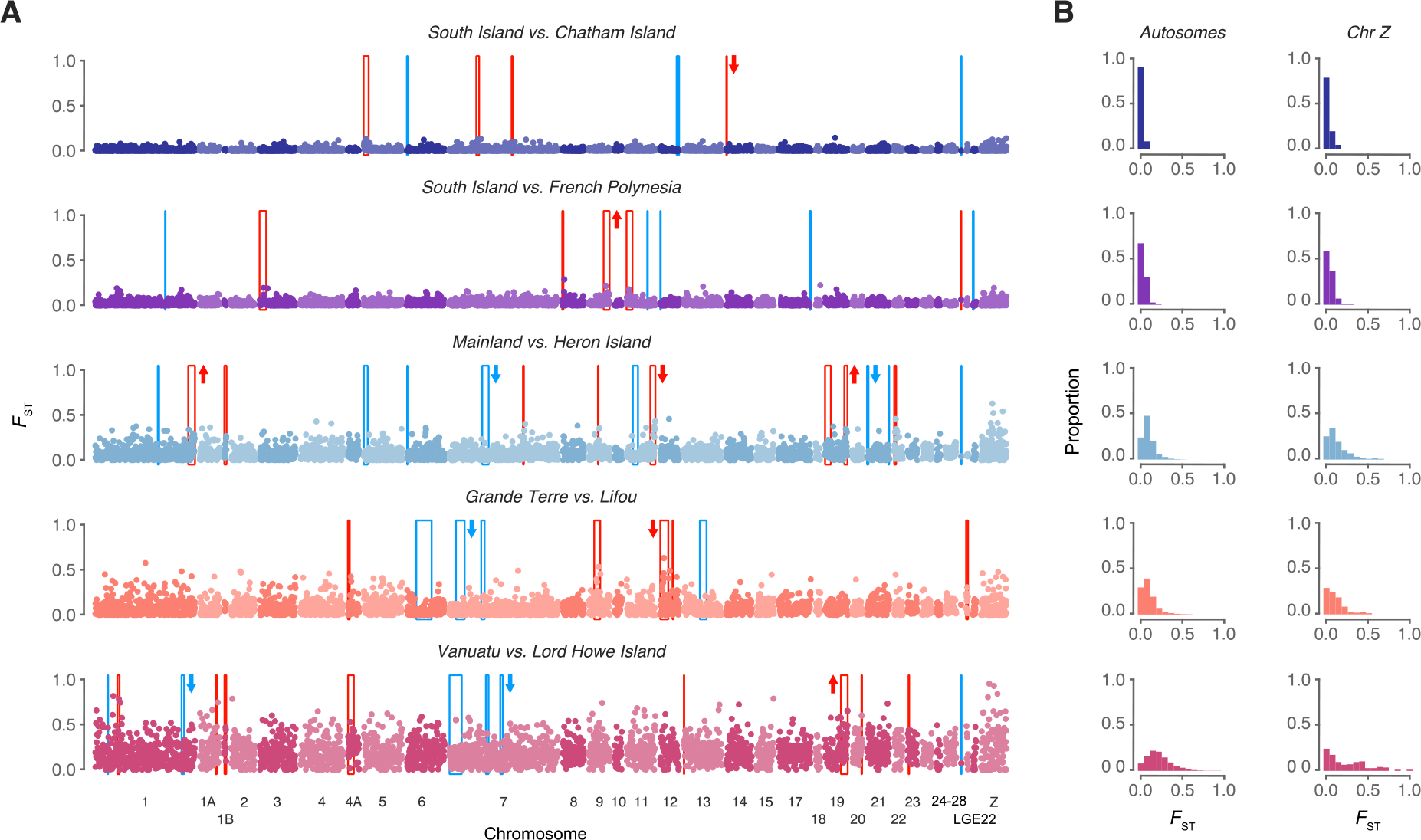
Distributions of divergence across the genome for diverging silvereye populations. (A)Pairwise *F*_ST_ across the genome for each population comparison calculated in non-overlapping 50kb windows. Regions of elevated differentiation (genomic islands) are highlighted in red and regions with low differentiation (genomic valleys) are highlighted in blue. Arrows beside genomic islands/valleys indicate if *d*_xy_ within these regions was significantly elevated (upwards pointing arrows) or decreased (downwards pointing arrows) compared to chromosomal background levels. Significant determined using Wilcoxon signed-rank tests. Chromosomes are numbered according to the zebra finch nomenclature. **(B)** Distributions of windowed *F*_ST_ values calculated in 50kb windows for each comparison. Distributions are shown for Autosomes and Chromosome Z separately.

**Table 2:** P-values for pairwise comparisons of *F*_ST_ distributional skew tested using a randomisation test. Below diagonal = autosomes only; above diagonal = chromosome Z only.

|  | **SI vs. CI** | **SI vs. FP** | **ML vs. HI** | **GT vs. LF** | **VN vs. LH** |
| --- | --- | --- | --- | --- | --- |
| **SI vs. CI** | - | 0.092 | 0.778 | 0.344 | 0.04 |
| **SI vs. FP** | 0.055 | - | 0.512 | 0.509 | 0.673 |
| **ML vs. HI** | 0.001 | 0.01 | - | 0.666 | 0.097 |
| **GT vs. LF** | 0.001 | 0.472 | 0.101 | - | 0.269 |
| **VN vs. LH** | 0.001 | 0.001 | 0.001 | 0.001 | - |

Like autosomes, chromosome Z showed a clear trend towards decreasing skew (Figure 3B), with divergence visually increasing over longer timeframes (Figure 4A). As was the case for autosomes, the *F*_ST_ distribution for Vanuatu vs. Lord Howe Island (late stage - no gene flow) was the least skewed, meaning that divergence was more widespread across the Z chromosome. However, this difference in skew was only significant when compared to the South Island vs. Chatham Island comparison (*P* <0.001, Table 2).

### Dynamics of genomic islands of divergence and valleys of similarity

Genomic islands were identified in all population comparisons, including populations diverging with gene flow (South Island vs. Chatham Island, Mainland vs. Heron Island, and Grande Terre vs. Lifou) as well as populations diverging in the absence of gene flow (South Island vs. French Polynesia and Vanuatu vs. Lord Howe Island) (Figure 4A). The number of genomic islands identified did not differ when compared across different gene flow scenarios or across different divergence timeframes. Similarly, genomic island size – although variable (mean size ranged from 110 kb to 1,490 kb, see Table 3 and Figure 5) – did not differ significantly across different gene flow contexts (Kruskal-Wallis: : 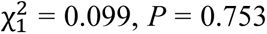) or divergence timescales (Kruskal-Wallis: : 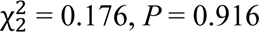). We identified four regions of the genome in which genomic islands of divergence occurred at the same location in two or more population comparisons. The size of genomic islands at shared locations is reported in Table 4. In most instances, absolute divergence (*d*_xy_) within individual genomic islands was not significantly different from background chromosomal levels. However, genomic islands with significantly elevated *d*_xy_ were identified in small numbers, as were genomic islands with below background level *d*_xy_ (Figure 4A).

**Figure 5:**
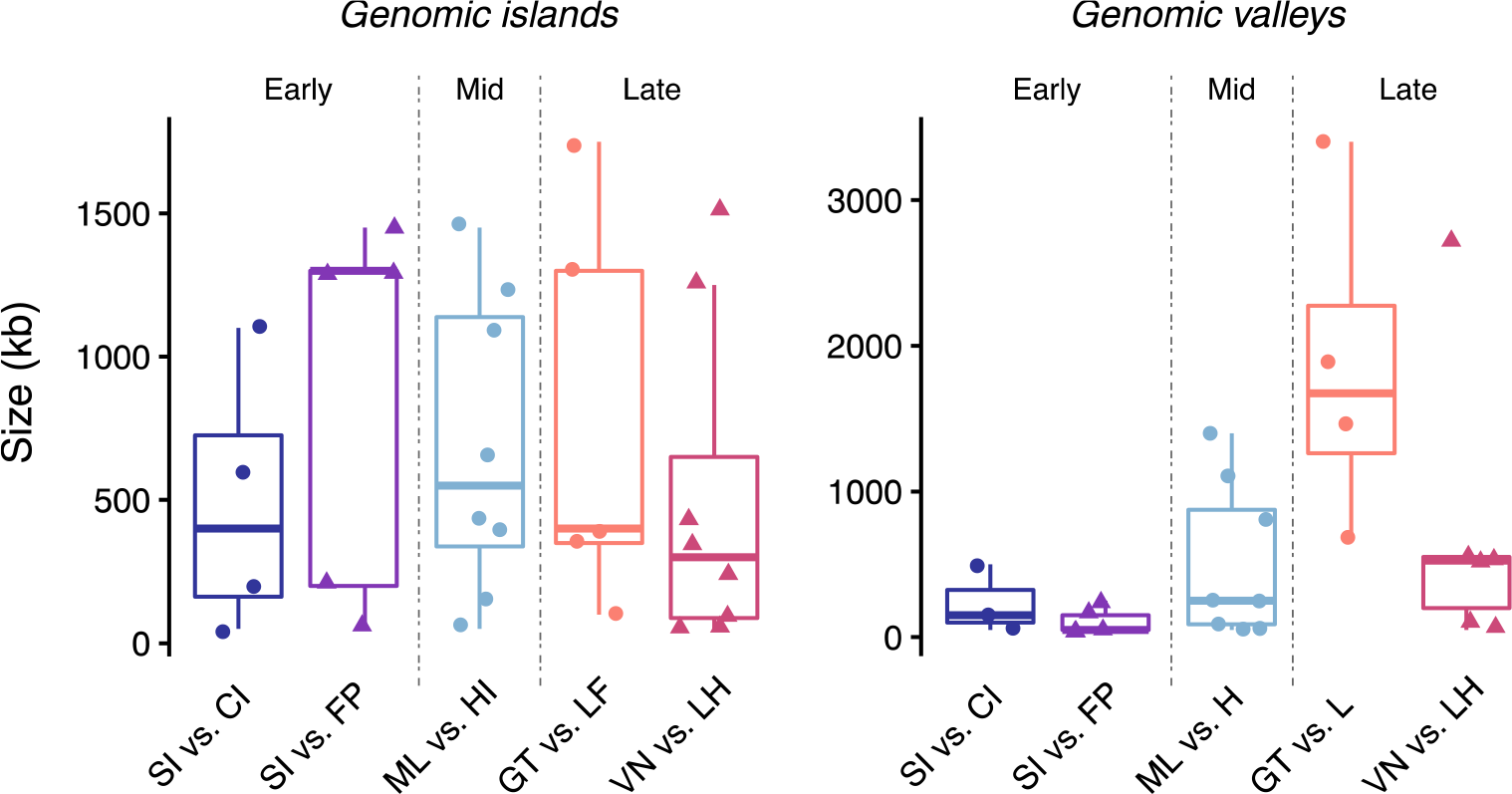
Size of genomic islands and genomic valleys identified for each population comparison. Populations diverging with gene flow are indicated by circles and populations diverging in isolation by triangles.

**Table 3:** Summary statistics of genomic islands and genomic valleys identified for each population comparison.

| Divergence<br>timeframe | Mode of<br>divergence | Comparison | No. identified | Mean size<br>(kb) | Max size<br>(kb) | Mean<br>$F_{ST}$ | Mean<br>$d_{xy}$ |
| --- | --- | --- | --- | --- | --- | --- | --- |
| **Genomic islands** |  |  |  |  |  |  |  |
| Early stage | Gene flow | SI vs. CI | 4 | 1490 | 5750 | 0.052 | 0.153 |
| Early stage | No-gene flow | SI vs. FP | 5 | 175 | 500 | 0.066 | 0.139 |
| Mid stage | Gene flow | ML vs. HI | 8 | 390 | 850 | 0.211 | 0.176 |
| Late stage | Gene flow | GT vs. LF | 5 | 110 | 200 | 0.288 | 0.136 |
| Late stage | No-gene flow | VN vs. LH | 8 | 341.7 | 750 | 0.385 | 0.220 |
| **Genomic valleys** |  |  |  |  |  |  |  |
| Early stage | Gene flow | SI vs. CI | 3 | 280 | 550 | 0.001 | 0.167 |
| Early stage | No-gene flow | SI vs. FP | 5 | 283.3 | 1050 | 0.003 | 0.162 |
| Mid stage | Gene flow | ML vs. HI | 8 | 205 | 750 | 0.023 | 0.131 |
| Late stage | Gene flow | GT vs. LF | 4 | 300 | 550 | 0.026 | 0.095 |
| Late stage | No-gene flow | VN vs. LH | 6 | 333.3 | 550 | 0.072 | 0.131 |

**Table 4:** Size of genomic islands and valleys occurring at the same location across multiple population comparisons.

| Divergence<br>timeframe | Mode of<br>divergence | Comparison | Genomic island<br>size (kb) |  |  |  | Genomic valley<br>size (kb) |  |  |  |
| --- | --- | --- | --- | --- | --- | --- | --- | --- | --- | --- |
|  |  |  | 1B | 4A | 9 | 19 | 6 | 7 #1 | 7 #2 | 27 |
| Early stage | Gene flow | SI vs. CI | - | - | - | - | 150 | - | - | 50 |
| Early stage | No-gene flow | SI vs. FP | - | - | - | - | - | - | - | - |
| Mid stage | Gene flow | ML vs. HI | 450 | - | 50 | 650 | 50 | - | 1400 | 50 |
| Late stage | Gene flow | GT vs. LF | - | 350 | 1300 | - | - | 1900 | 700 | - |
| Late stage | No-gene flow | VN vs. LH | 350 | 1250 | - | 1500 | - | 2700 | 550 | 50 |

Genomic valleys were identified in all diverging population comparisons (Figure 4A). However, the number identified, ranging from 3 to 10 (Table 3), was not related to divergence timescale or gene flow context (all Kruskal-Wallis *P*-values > 0.05). Mean genomic valley size, was highly variable (Figure 5). Genomic valley size differed significantly when compared across divergence timeframes (Kruskal-Wallis: 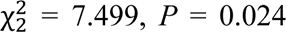) but not gene flow scenarios (Kruskal-Wallis: 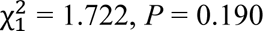)). We identified four regions of the genome in which a genomic valley occurred across multiple population comparisons (Table 4). In most instances, absolute divergence (*d*_xy_) within individual genomic valleys was not significantly different from background chromosomal levels. However, genomic valleys with significantly decreased *d*_xy_ were identified in small numbers (Figure 4A). Genomic valleys with significantly elevated *d*_xy_ were not identified.

### Correspondence of genomic islands and valleys to the position of outlier SNPs

*PCAdapt* identified 1 to 330 outlier SNPs in each population comparison (Table 5 and Figure 6). For the late stage – no gene flow comparison (Vanuatu vs. Lord Howe Island), 4 outlier SNPs fell within genomic islands and 3 outliers fell within genomic valleys. For the remaining comparisons, outlier SNPs did not correspond to the position of genomic islands or genomic valleys (Table 5 and Figure 6).

**Figure 6:**
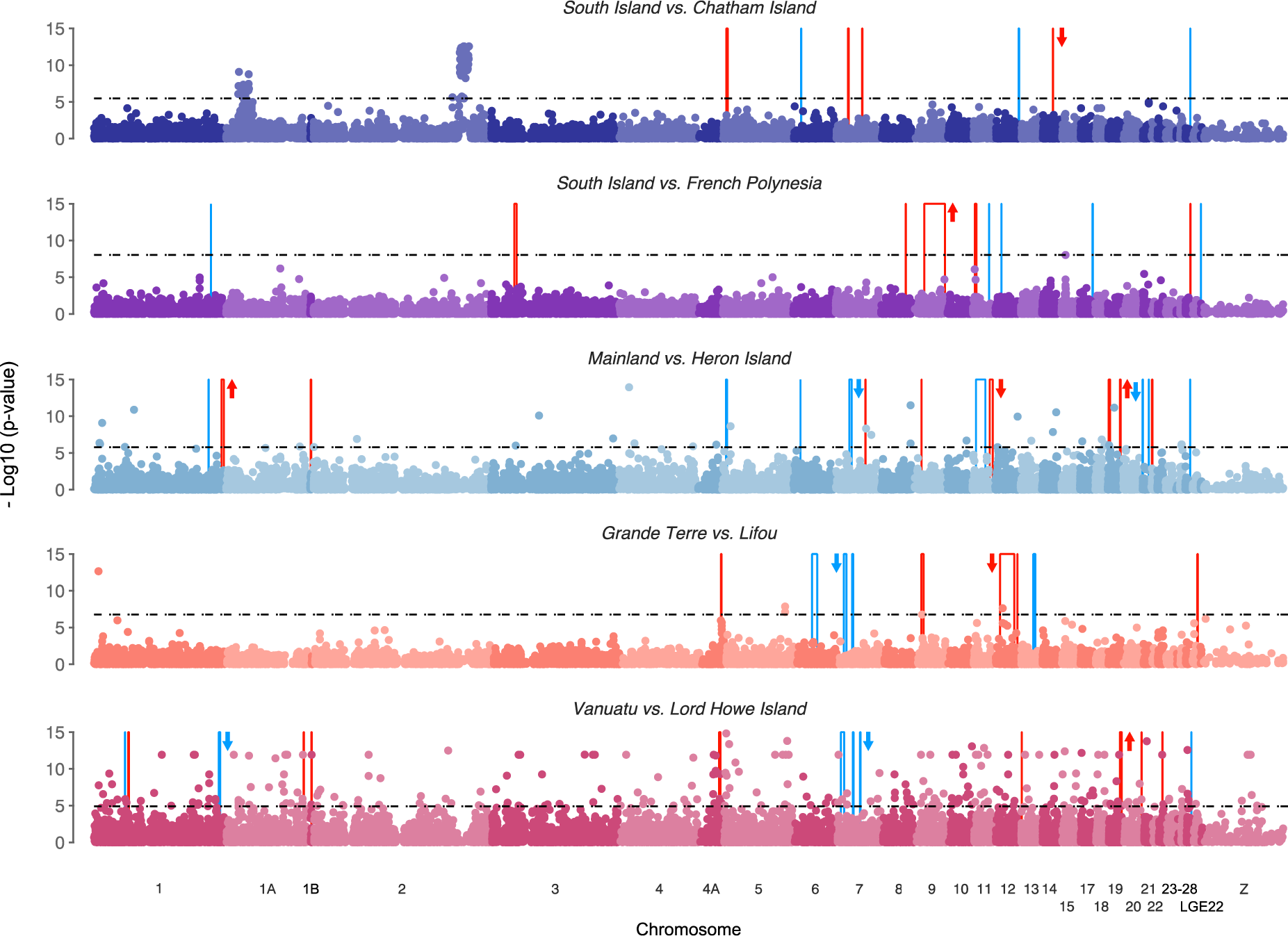
Manhattan plot of negative log10 (P values) estimated using PCAdapt. Points above the dashed line indicate outlier SNPs identified using FDR = 0.001. Highlighted points indicate outlier SNPs identified using FDR = 0.001. 50kb windows with elevated differentiation (genomic islands) are highlighted in red and regions with low differentiation (genomic valleys) are highlighted in blue. Genomic islands and genomic valleys are based on *F*_ST_ and locations are the same as those shown in Figure 4. Arrows beside genomic islands/valleys indicate if dxy within these regions was significantly elevated (upwards pointing arrows) or decreased (downwards pointing arrows) compared to chromosomal background levels. Significant determined using Wilcoxon signed-rank tests. Chromosomes are numbered according to the zebra finch nomenclature.

**Table 5:**
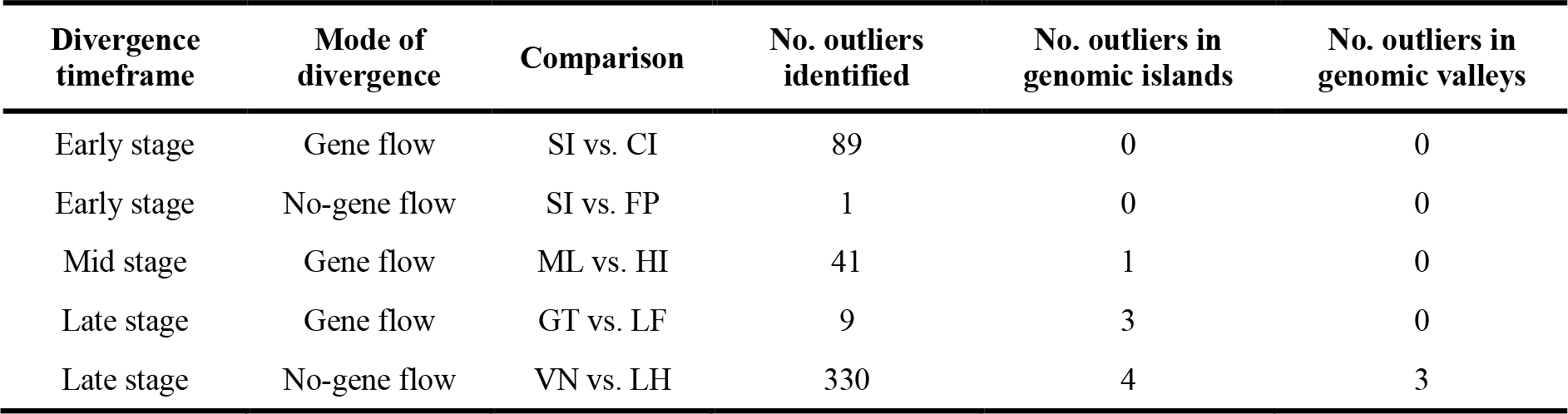
Outlier SNP summaries. Number of outlier SNPs identified using *PCAdapt* and the number located within genomic islands and valleys per population comparison.

### Candidate gene analysis

A review of the literature identified 22 candidate genes associated with body and/or beak size differences in birds, for which we had sequence data or occurred within the 50kb windows used when summarising divergence statistics (Table 6). Of these, four (*BMPR1A*, *COL4A5*, *ITPR2* and *VPS13B*) contained outlier SNPs in at least one population comparison. For the late stage – no gene flow comparison (Vanuatu vs. Lord Howe Island) the genomic island located on chromosome 4A contained *BMPR1A*.

**Table 6:**
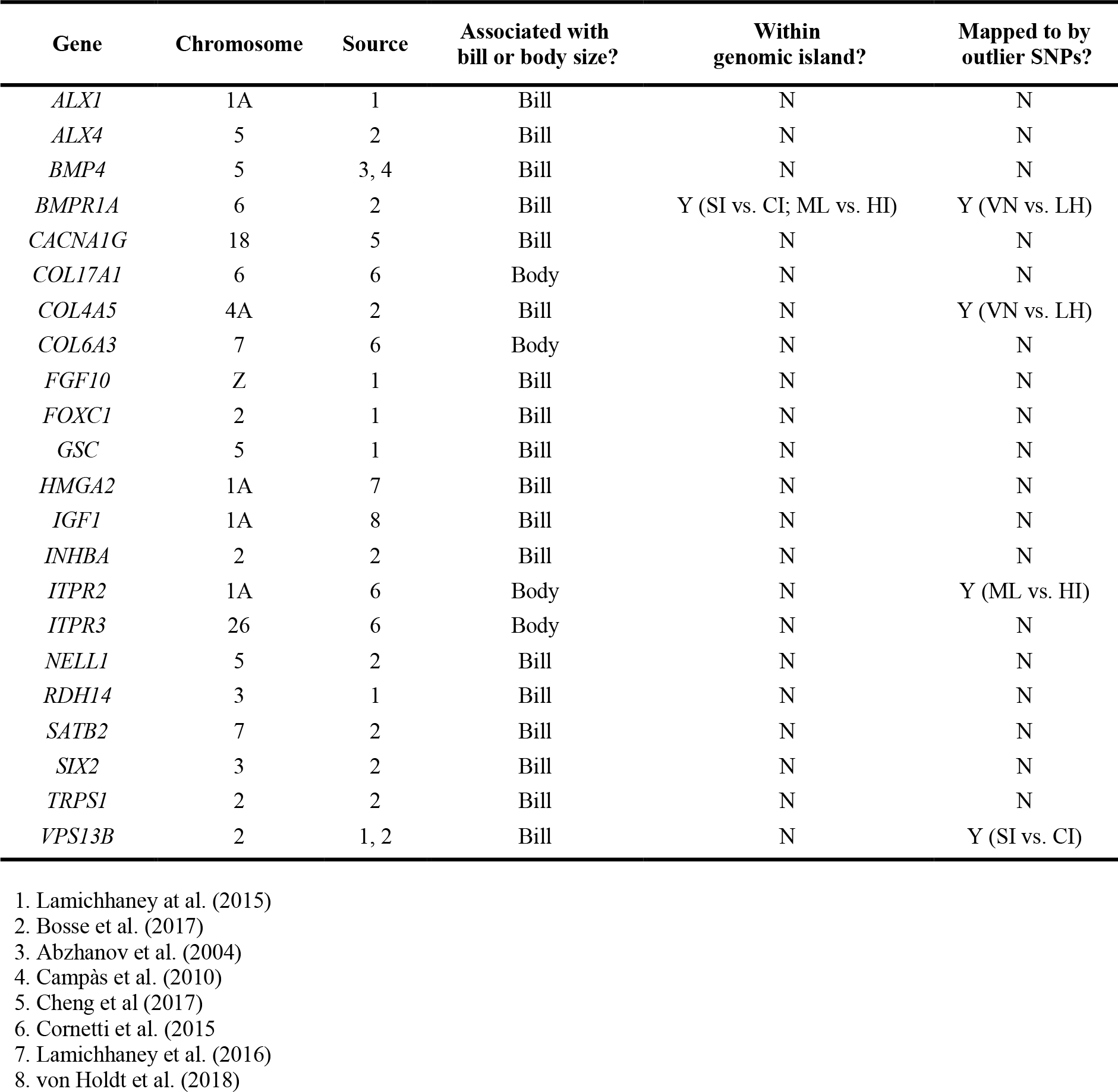
Patterns of differentiation in genes linked to body/bill size differentiation in birds.

### Simulations of neutral divergence

Simulations of neutral divergence, using an individual based model, were able to capture the transition from localised to more genome-wide levels of divergence, in which *F*_ST_ distributions were characterised by low values of *F*_ST_ during the early stages of divergence, shifting towards higher *F*_ST_ values over longer divergence timeframes (Figure 7). However, differences in distributional skew between simulations at different timescales were not all significant (Table 7). Simulating divergence under various levels of gene flow (*m* = 0, *m* = 0.0001, *m* = 0.001, *m* = 0.01) showed that the accumulation of divergence slowed under increasing levels of gene flow (Figure 8).

**Figure 7:**
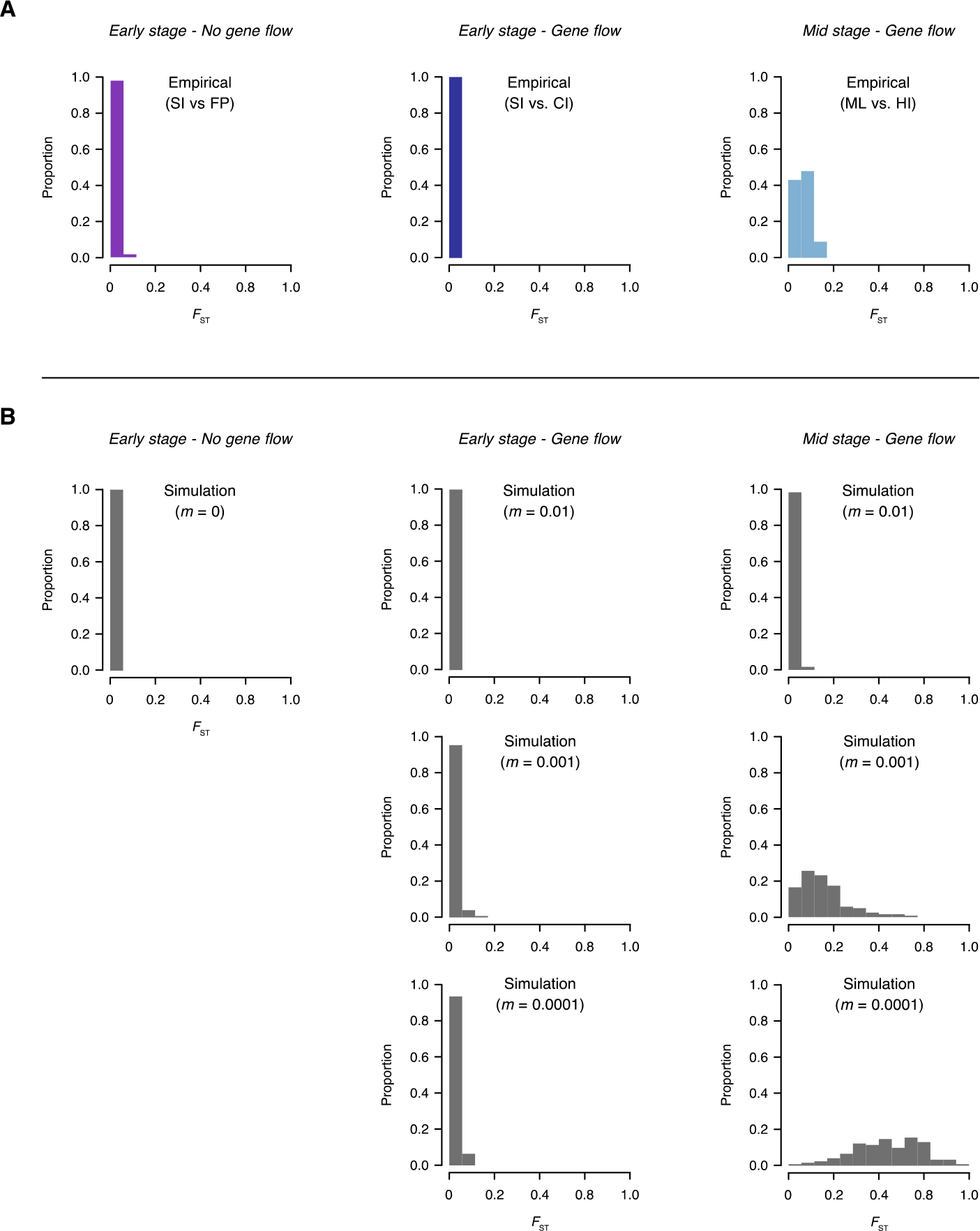
Comparison of observed and simulated *F*_ST_ distributions. (A) Frequency distributions of pairwise differentiation (*F*_ST_) for empirical data (chromosome 5 only). (B) Frequency distributions of pairwise differentiation (*F*_ST_) for simulated data. The *F*_ST_ values are calculated in 500kb windows. Simulations were conducted using a recombination rate of 3 cM/Mb (the distance between loci for which the expected average number of intervening chromosomal crossovers in a single generation is 0.01). Simulations of divergence with gene flow were conducted under different migration rates (*m* = 0.01, *m* = 0.001, and *m* = 0.0001). Simulation timeframes matched that for each empirical comparison (Early stage - no gene flow: 30 generations; Early stage - gene flow: 60 generations; Mid stage - gene flow: 1,000 generations).

**Figure 8:**
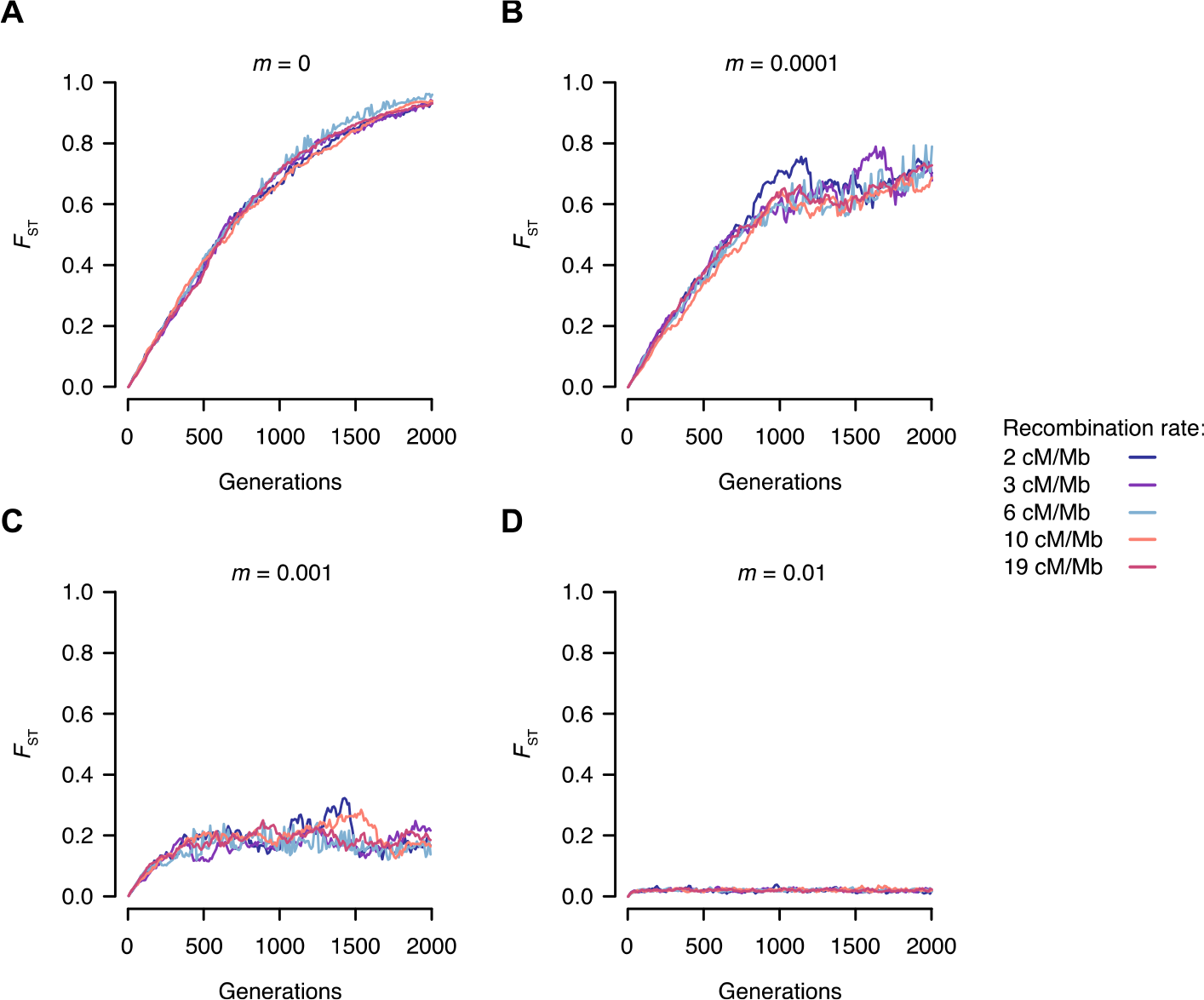
Expected *F*_ST_ values for 2,000 generations of simulated neutral divergence. (A) During divergence without gene flow (*m* = 0) *F*_ST_ rapidly increases, achieving fixation at 2,000 generations under all recombination rates; (B) During divergence with low levels of migration (*m* = 0.0001) *F*_ST_ increases rapidly, approaching fixation at 2,000 generations for all recombination rates; (C) During divergence with intermediate levels of migration (*m* = 0.001) *F*_ST_ increases rapidly during the initial ∼300 generations, after which it continues to fluctuate around 0.2 with some variation between recombination rates; and (D) During divergence with high levels of gene flow (m = 0.01) low levels of *F*_ST_ are maintained across all 2,000 generations of divergence.

**Table 7:** P-values for pairwise comparisons of *F*_ST_ distributional skew for simulations, tested using a randomisation test. Values in grey shaded cells indicate level of skew for a given timeframe.

| Gene flow | Simulation timeframe |  |  |  |
| --- | --- | --- | --- | --- |
|  | Generations | 30 | 60 | 1000 |
| m = 0 | 30 | 1.094 | 0.025 | 0.001 |
|  | 60 | - | 0.473 | 0.021 |
|  | 1000 | - | - | -0.064 |
| m = 0.0001 | 30 | 1.170 | 0.001 | 0.001 |
|  | 60 | - | 0.001 | 0.081 |
|  | 1000 | - | - | -0.352 |
| m = 0.001 | 30 | 0.593 | 0.386 | 0.103 |
|  | 60 | - | 1.983 | 0.079 |
|  | 1000 | - | - | 1.235 |
| m = 0.01 | 30 | 0.936 | 0.627 | 0.511 |
|  | 60 | - | 1.730 | 0.875 |
|  | 1000 | - | - | 1.830 |

## DISCUSSION

We take advantage of an exceptional natural system where gene flow scenarios and divergence timescales are known *a priori*, to reveal key insights into how divergence accumulates at the genomic level. Our results provide an empirical characterisation of the change in the distribution of genomic divergence using a proxy for the speciation continuum. The pattern of decreasing skew in the distribution of genomic divergence is consistent with the intuitive explanation of how heterogeneity originates, and divergence accumulates, i.e. divergence at a small number of selected loci followed by divergence hitchhiking. However closer inspection of the genomic landscape provides equivocal support for this particular mechanism. Genomic islands were rarely associated with SNPs putatively under selection and genomic islands did not widen as expected under the divergence hitchhiking model of speciation. Genomic valleys of similarity were also a common feature of the heterogeneous genomic landscape, but we did not find clear evidence to attribute these to selective processes. Furthermore, by simulating divergence under a neutral model we were able to capture the transition from localised to more genome-wide levels of divergence. The transition from localised divergence (highly skewed distribution) to more genome-wide levels of divergence (a more even distribution of genetic differentiation) therefore appears to occur largely outside of the context of genomic islands and is not dependent of selection, raising questions about the divergence hitchhiking model of speciation, where divergence is initiated through selection of a small number of genes (presumably of large effect), genomic islands form around them, and divergence hitchhiking completes genome-wide divergence.

### The accumulation of divergence across the genome

Our results demonstrate how divergence accumulates across the genome as populations move along the speciation continuum. *Zosterops lateralis* populations at the earliest stages of divergence followed extreme L-shaped *F*_ST_ frequency distributions where divergence was limited to few loci, with a reduction of skew in *F*_ST_ values as divergence became more wide-spread at later stages of the speciation continuum. This shift in distribution occurred in both the presence and absence of gene flow. We are aware of only two other studies where similar comparisons could be made: *Heliconius* butterflies (Martin et al., 2013) and *Ficedula* flycatchers (Burri et al., 2015). Unlike Burri et al. (2015), we did not observe a shift towards an extreme left-skewed distribution where the majority of SNPs reach fixation. This is likely due to the inclusion of comparisons between flycatcher species with divergence times of millions of years, whereas our within-species divergence timeframes were too short for an extreme left-skewed distribution to be reached.

On the whole however, the observed distributions of genomic differences across the genome fit with expectations for how genomic divergence accumulates across the speciation continuum (Nosil and Feder, 2012; Seehausen et al., 2014). However, our simulations demonstrated that the shift from localised to more genome-wide levels of divergence could be achieved assuming a neutral model of molecular evolution. This finding was robust to varying levels of recombination, which is interesting given the emphasis on linkage disequilibrium in driving localised and ultimately genome-wide divergence (Seehausen et al., 2014). One important difference, however, was that our simulations suggest that the transition from localised to more genome-wide levels of divergence occurs at a faster rate than is observed empirically - under allopatry and low-level gene flow *F*_ST_ approached fixation by 2,000 generations. This suggests that some other mechanism, such as frequency-dependent selection, may be maintaining genetic variation at individual SNPs.

When matched for divergence timeframe, the pace at which divergence accumulated across the genome was moderated by gene flow. This was manifested as higher distributional skew of autosomal *F*_ST_ values for silvereye populations diverging under gene flow compared to populations diverging in isolation. Martin et al. (2013) found that genomic divergence was more widespread for races and species of *Heliconius* butterflies diverging in geographic allopatry than for those diverging in parapatry and sympatry, suggesting an underlying role for gene flow in slowing the accumulation of genomic divergence. We provide a further empirical characterisation of this difference in the accumulation of genomic divergence. This moderating effect of gene flow was also apparent when simulating divergence.

Given that the silvereye population in French Polynesia was the product of a human introduction and therefore likely had lower effective founder size than natural colonisations involving large expanding populations of silvereyes, we cannot rule out that the more wide-spread genomic divergence observed for the South Island vs. French Polynesia population comparison (when compared to other early stage comparison: South Island vs. Chatham Island) is the product of a potential severe population bottleneck during introduction. The potential for population founding to accelerate the accumulation of genomic differences has been highlighted previously (Ravinet et al., 2017), and in the silvereye system could be addressed through genomic comparisons of a number of other populations with varying intensities of population bottlenecks (Estoup and Clegg, 2003).

Divergence of silvereye chromosome Z was elevated beyond that observed for autosomes, as seen in a number of other avian divergence studies (Storchová et al., 2010; Sætre and Sæether, 2010; Ruegg et al., 2014; Oyler-McCance et al., 2015). Sex chromosomes are thought to contain a large number of genes related to sexual selection and reproductive isolation, and as such have been proposed to play a disproportionately large role in speciation (Dobzhansky, 1974; Coyne, 1985; Ellegren, 2009; Charlesworth et al., 2017). Despite this, no genomic islands, and only a handful of significant outlier SNPs (in the Vanuatu vs. Lord Howe Island comparison only), were identified within chromosome Z. Instead, the elevated levels of divergence may be partly explained by the lower effective population size of the Z chromosome (3/4 of an autosome), amplifying the effects of genetic drift (Mank et al., 2010).

### The dynamics of genomic islands of divergence and valleys of similarity

Our results run counter to several patterns expected from the divergence hitchhiking model of speciation. First, genomic islands of divergence were seen in similar frequency in populations diverging both with and without gene flow. This adds to a number of empirical studies in natural systems where genomic islands have been observed in populations that are, or are assumed to be, genetically isolated (Renaut et al., 2013; Marques et al., 2016; Han et al., 2017; Van Doren et al., 2017; Zhang et al., 2017). Second, genomic islands were not wider on average for older divergence timeframes, which does not align with the prediction that genomic islands expand with time as linkage disequilibrium facilitates divergence of neutral and weakly selected loci via divergence hitchhiking (Smadja et al., 2008; Via and West, 2008; Via, 2012). A small number of recent studies have also failed to find a positive association between divergence time and average genomic island size (*Timema* stick insects: (Riesch et al., 2017); Darwin’s finches: (Han et al., 2017); and Yellow-rumped warblers: (Toews et al., 2016)). Taken together, growth of genomic islands may not be pervasive in the generation of genome-wide divergence, with the accumulation of divergence in our example instead taking place largely outside of the context of genomic islands. This observation is consistent with the more recently proposed multi-locus coupling mechanism of divergence (Flaxman et al., 2013, 2014; Butlin and Smadja, 2017; Schilling et al., 2018).

Associating average genomic island size with divergence time may be too coarse an approach, and comparisons of genomic islands at specific genomic locations may be a better method to understand how time and gene flow shape growth of genomic islands. In our case, only four genomic islands were located at overlapping genomic positions across population comparison. Of the three that occurred across different timeframes, two were largest in the late stage comparison and the other smallest. Whole genome analysis would likely provide more shared genomic islands to allow statistical comparisons, but currently we have equivocal support for the idea that expansion of genomic islands of divergence at specific locations over time regularly occurs.

Contrary to the expectation that genomic islands of divergence contain loci under strong divergent selection (Wu, 2001), we find that genomic islands regularly occur in regions of the *Z. lateralis* genome where outlier SNPs putatively under directional selection were not identified. A similar result was reported in diverging Swainson’s thrush (*Catharus ustulatus*) subspecies (Ruegg et al., 2014). Hence, it appears that genomic islands of divergence are not always seeded by directional selection, but whether this is frequently the case requires a broader number of empirical examples. The formation of genomic islands without obvious evidence of directional selection highlights the need for alternative explanations for their formation (Turner et al., 2005; Hohenlohe et al., 2010; Feder et al., 2013). One such explanation is that genomic islands of divergence may arise in regions of low recombination such as centromeres (Carneiro et al., 2009; Renaut et al., 2013; Burri et al., 2015). However, lack of recombination and karyotype maps for silvereyes at present prevents formal testing of this hypothesis here. In addition to variation in recombination rate, other genomic features that may contribute to the formation of genomic islands include variation in gene density and variation in mutation rate along the genome (Ravinet et al., 2017). Whole genome sequencing would help to address the contribution of these genomic features.

A comparison of absolute and relative measures of divergence (e.g. *d*_xy_ and *F*_ST_ respectively) can provide further insight into the formation of genomic islands (Nachman and Payseur, 2012; Han et al., 2017; Delmore et al., 2018). Unlike *F*_ST_, which is consistently elevated around loci under directional selection, *d*_xy_ may be elevated, reduced or unchanged. Under divergence with gene flow, genomic islands with elevated measures of both relative and absolute divergence are expected to form around regions where gene flow is disadvantageous, such as around regions containing variants involved in local adaptation or reproductive isolation (Han et al., 2017). We observed such a pattern in only one of the three comparisons diverging with gene flow. However, as we also observed islands where both types of measures were elevated in populations diverging without gene flow, our results provide support for an alternative explanation for this pattern: that genomic islands with elevated *d*_xy_ occur due to lineage sorting (Han et al., 2017). In the absence of gene flow, selective sweeps, which convert between-population variation into fixed differences, are expected to produce regions of elevated *F*_ST_ but not *d*_xy_ (Nachman and Payseur, 2012). Such islands were the most frequently observed across all comparisons, suggesting that genomic islands arise most frequently within regions where gene flow is absent. Genomic islands (based on *F*_ST_) but with reduced *d*_xy_, were the least frequently observed. Such a pattern is thought to be caused by background selection and recurrent selective sweeps, both of which result in locally reduced levels of genetic variation (Seehausen et al., 2014; Wolf and Ellegren, 2016; Ma et al., 2017).

Genomic valleys play an important role in shaping the genomic landscape across the speciation continuum, occurring across all population comparisons. In particular, their occurrence during the late stage of divergence may aid maintenance of heterogeneity by slowing the approach to genome-wide divergence at some genomic regions. Should genomic valleys remain present in the very advanced stages of divergence - i.e. comparisons at the species level - these regions likely contribute to the formation of the tail of extreme values that characterises the left skewed distributions observed during the late stage of divergence such as those observed between diverging *Ficedula* flycatcher species by Burri et al. (2015). Comparisons between *Zosterops* species are needed to confirm this expectation.

A comparison of the nature of genomic valleys across the speciation continuum and gene flow contexts allows us to comment on mechanisms proposed to generate genomic valleys. Although genomic valleys frequently showed a pattern of reduced *F*_ST_ and *d*_xy_, as would be expected if genomic valleys form at loci under parallel selection (Roesti et al., 2014), the position of outlier SNPs corresponded to the position of genomic valleys in only a single comparison (Vanuatu vs. Lord Howe Island). As such, at present we have limited evidence to confirm that genomic valleys frequently arise due to effects of parallel selection. Second, as the number of genomic valleys was found to be stable over time and their size was found to be overall larger in populations diverging over longer timescales, we do not find support that genomic valleys are solely the product of incomplete lineage sorting at neutral loci, as under this mechanism genomic valleys would be expected to become smaller and less numerous as shared ancestral variation breaks down over time (Stölting et al., 2013).

### Repeated evolutionary patterns

While we found limited evidence that candidate genes thought to be associated with body and/or bill size differences in passerines were concentrated within genomic islands, only one genomic island contained a candidate gene *BMPR1A*, we did find evidence that such genes may be implicated in the repeated evolutionary pattern seen in island silvereyes. Four genes (*BMPR1A*, *COL4A5*, *ITPR2* and *VPS13B*) are potentially under selection in the Vanuatu vs. Lord Howe Island, Mainland vs. Heron Island and South Island vs. Chatham Island comparisons respectively. Outliers SNPs identified for these comparisons mapped to these specific genes and for the Vanuatu vs. Lord Howe Island comparison *COL4A5* also contained SNPs with fixed differences. *COL4A5* is a type IV collagen gene associated with craniofacial disease in humans (Jonsson et al., 1998), and more recently associated with bill length in the great tit (Parus major) (Bosse et al., 2017). *ITPR2* plays a key role in metabolism and growth (Cornetti et al., 2015) and *VPS13B* has been associated with bill morphology in Darwin’s finches (Lamichhaney et al., 2015) and facial dysmorphism in humans (Balikova et al., 2009).

## CONCLUSION

While the goal of identifying unambiguous signatures of particular evolutionary processes in patterns of genomic divergence remains elusive, the addition of empirical studies of well-characterised systems provides valuable insight into the nature of divergence across the speciation continuum. In the silvereye system, gene flow is clearly important in shaping genome-wide divergence. While these results provide an excellent baseline for inferring the role of gene flow in other silvereye populations, equivalent calibrations would likely be needed on a species-by-species basis for the role of gene flow in a population of unknown history to be assessed. Genome-wide divergence in silvereyes does not hinge on the formation and growth of genomic islands. This is at odds with the intuitive and until recently, frequently invoked verbal model describing the accumulation of genomic divergence during the speciation continuum. Instead, differences are spread in a remarkably even way across the genome, a finding that is perhaps more in tune with the mutli-locus coupling model of divergence. The joint empirical and theoretical approach we present offers a potentially powerful tool to test a range of hypotheses about the mechanisms that underlie genomic speciation.

## ACKNOWLEDGEMENTS

This work was funded by a grant from the John Fell Oxford University Press Research Fund to T.C. and S.M.C., with additional support from an anonymous donation to K.C.R. and a Natural Environment Research Council (NERC) studentship awarded to A.T.S.P. C.S.Q. acknowledges support from the Early Postdoc Mobility fellowship of the Swiss National Science Foundation (P2GEP3_168973). Sample collection was conducted with the permission of the governments of Australia, French Polynesia, New Caledonia, New Zealand and Vanuatu. We thank the Museum d’Histoire Naturelle de Genève, Alice Cibois, Nick Clark, Rob Fleischer, and Ally Phillimore for providing additional samples. The authors would like to acknowledge the use of the University of Oxford Advanced Research Computing (ARC) facility in carrying out this work.

## AUTHOR CONTRIBUTIONS

S.M.C., K.C.R, E.C.A and T.C. conceived the project. S.M.C. and K.C.R supervised the project. V.L.U. conducted the laboratory work. Bioinformatics and analyses were primarily carried out by A.T.S.P. with assistance from S.M.C., K.C.R, E.C.A., B.V.D. and T.C. Simulations were conducted by C.S.Q with input from T.C. and S.M.C. The manuscript was written by A.T.S.P., S.M.C., and K.C.R. with input on individual sections by T. C., and C.S.Q. and editing by all authors.

## DATA ACCESS

Resequencing data from this study have been submitted to the National Center for Biotechnology Information (NCBI; https://www.ncbi.nlm.nih.gov) under accession number PRJNA489169.

