## Supplementary Materials for "The genomic landscape of divergence across the speciation continuum in island-colonising silvereyes (*Zosterops lateralis*)"

### SUPPLEMENTARY METHODS

#### *Library preparation*

100 ng of genomic DNA from each individual was digested with 2.4 U of restriction enzyme SbfI-HF (New-England Biolabs Inc., Beverly MA, USA) at $37_{o}$C for 60 minutes, followed by an inactivation step of $80_{o}$C for 20 minutes. BestRAD Sbf1 adapters were then ligated to the overhanging ends of the products of the restriction reactions by adding 50 nM to each sample with 320 U T4 DNA Ligase (New-England Biolabs Inc.). Barcoding of samples was achieved with a set of index nucleotides within the BestRAD Sbf1 adapter sequences. Reactions were incubated at $20_{o}$C for 16 hours and then heat-inactivated by holding at 65°C for 20 minutes. The reactions were then pooled, cleaned with AMPure XP beads, and the products randomly sheared to a mean size of 500 bp by sonication, using a BioRuptor NGS (Diagenode). Following sonication, Dynabeads M-280 streptavidin magnetic beads (Life Technologies) were used to bind the biotinylated ends of the BestRAD Sbf1 adaptors. Sbf1-HF was used once more to release adapter bound fragments and AMPure XP beads were used to clean the samples. After the bead clean up, ends were treated with Blunt End-Repair Mix found in the NEBNext Ultra DNA Library Prep Kit (New-England Biolabs Inc.) to remove overhangs. NEBNext Adaptors for Illumina sequencing were then ligated to blunt ends following a two-step incubation; first incubated at $20_{o}$C for 15 minutes with Blunt/TA Ligase Master Mix (New-England Biolabs Inc.) and then at $37_{o}$C for 15 minutes with USER Enzyme (New-England Biolabs Inc.). Following size selection using AMPure XP beads to isolate fragments within the size range 300–700 bp, the library was PCR enriched using 10 uM of P1 index primer and 10 uM of Universal PCR Primer (NEB). First a test PCR was run using 5 ul of DNA with 15 PCR cycles. Depending on the brightness of the band produced, the final PCR cycle number was adjusted from 9-12 cycles. PCR products were cleaned using AMPure XP beads. Libraries were sequenced on three Illumina HiSeq4000 lanes (Illumina, San Diego, CA, USA) at the UC Davis Genome Center using paired-end 150-bp sequence reads.

### SUPPLEMENTARY TABLES & FIGURES

**Table S1. List of parameters of the model with default values**

| Parameter | Definition | Value | Source |
|---|---|---|---|
| $N_0$ | Initial number of breeders | 400 | Kikkawa & Wilson (1983) |
| $\zeta$ | Number of young fledging by clutch | 1.9 | Kikkawa & Wilson (1983) |
| $S_S$ | Survival rate at summer | 0.96 | Sandvig et al. (2017) |
| $S_a$ | Survival rate at autumn | 0.76 | Sandvig et al. (2017) |
| $S_w$ | Survival rate at winter | 0.63 | Sandvig et al. (2017) |
| $\delta$ | Sex ratio | 0.5 | Kikkawa & Wilson (1983) |
| $G$ | Generation time | 3 | Clegg et al. (2008) |
| $n_L$ | Number of SNPs | 2533 | Chromosome 17 |
| $\sigma_{dem}$ | Stochastic demographic variant | 0.0013 | * |
| $\theta$ | Recombination rate | 2, 3, 6, 10, 19 | † |
| $m$ | Migration rate | 0.0001, 0.001, 0.01 | ‡ |

* This value gives an average of 200 breeding pairs per generation, in line with the size of the breeding population observed on Heron Island (Kikkawa and Wilson 1983).

† Recombination rates observed for parts of the collared-flycatcher (*Ficedula albicollis*) genome (1). The value of 2 cM/Mb is the average for chromosomes > 100Mb, 3 cM/Mb is the genome-wide average; 6 cM/Mb the average for chromosome 17, 10 cM/Mb the average for chromosomes < 10Mb, and 19 cM/Mb the maximum average observed in a collared-flycatcher chromosome.

‡ A migration rate of 0.01 was empirically estimated among Heron Island and neighbouring islands (Brook and Kikkawa 1998). This was taken as the upper estimate of migration rate, but as the population comparisons considered here include more isolated islands, we also ran simulations assuming migration rates one and two magnitudes lower.

**Table S2:** Distributional skew of $F_{ST}$ values calculated for each autosomal chromosome. Only chromosomes with at least 10 x 50kb windows are reported. Comparisons where distributional skew is significantly higher when diverging with gene flow are highlighted in grey. Significance determined using a modified randomisation test.

| **Chromosome** | **Early Stage** | | **Mid stage** | **Late stage** | |
|---|---|---|---|---|---|
| | Gene flow (SI vs. CI) | No gene flow (SI vs. FP) | Gene flow (ML vs. HI) | Gene flow (GT vs. LF) | No gene flow (VN vs. LH) |
| 1 | 2.19 | 1.93 | 1.52 | 2.02 | 1.36 |
| 1A | 1.54 | 1.38 | 1.79 | 1.88 | 1.01 |
| 1B | - | - | - | - | - |
| 2 | 3.13 | 1.11 | 1 | 2.31 | 1.14 |
| 3 | 2.32 | 2.3 | 1.56 | 1.61 | 0.9 |
| 4 | 2.66 | 1.72 | 1.58 | 1.94 | 0.95 |
| 4A | 1.24 | 1.68 | 1.14 | 1.73 | -0.14 |
| 5 | 3.04 | 1.37 | 1.75 | 1.36 | 0.92 |
| 6 | 3.17 | 1.6 | 1.51 | 1.60 | 0.66 |
| 7 | 2.99 | 1.56 | 1.55 | 2.07 | 0.97 |
| 8 | 1.91 | 3.28 | 1.7 | 0.77 | 0.84 |
| 9 | 1.25 | 2.41 | 1.83 | 1.61 | 1.1 |
| 10 | 1.28 | 0.78 | 0.23 | 0.82 | 0.59 |
| 11 | 2.23 | 1.8 | 2.22 | 1.76 | 1.2 |
| 12 | 1.72 | 0.93 | 2.23 | 1.66 | 0.56 |
| 13 | 2.82 | 2.32 | 1.13 | 1.34 | 1.22 |
| 14 | 1.64 | 1.2 | 1.01 | 1.32 | 1.01 |
| 15 | 1.32 | 1.28 | 1.76 | 1.93 | 1.33 |
| 17 | 1.79 | 1.73 | 1.32 | 1.60 | 0.76 |
| 18 | 1.99 | 3.26 | 0.49 | 0.82 | 0.64 |
| 19 | 4.37 | 1.91 | 1.16 | 0.52 | 0.97 |
| 20 | 1.45 | 1.82 | 2.05 | 2.53 | 0.59 |
| 21 | 2.06 | 0.89 | 1.76 | 1.05 | 1.06 |
| 22 | 1.72 | 0.81 | 1.71 | 2.21 | -0.02 |
| 23 | 1.3 | 0.56 | 1.57 | 1.89 | -0.06 |
| 24 | 1.56 | 0.92 | 0.38 | 0.99 | 0.69 |
| 25 | 1.54 | 1.28 | 1.68 | 0.16 | 1.09 |
| 26 | 2.33 | 0.25 | 0.64 | 0.79 | 0.46 |
| 27 | - | - | - | - | - |
| 28 | 1.36 | 1.03 | 0.15 | -0.04 | 0.41 |
| LGE22 | 2.21 | 0.9 | 1.08 | -0.17 | 0.74 |

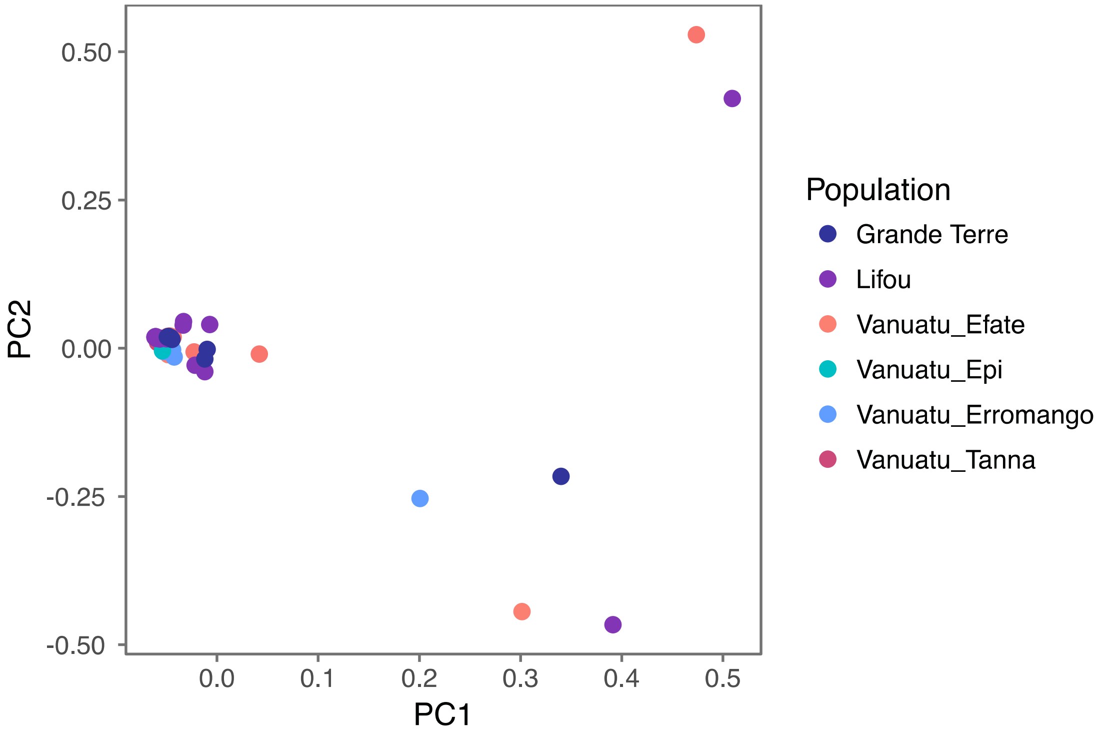


**Figure S1: Principle component analysis of genetic variation.** Based on 129,505 SNPs across 56 individuals sampled across 6 Melanesian islands (New Caledonia and Vanuatu). The variance explained by PC1 and PC2 is 16.46% and 10.47%, respectively.

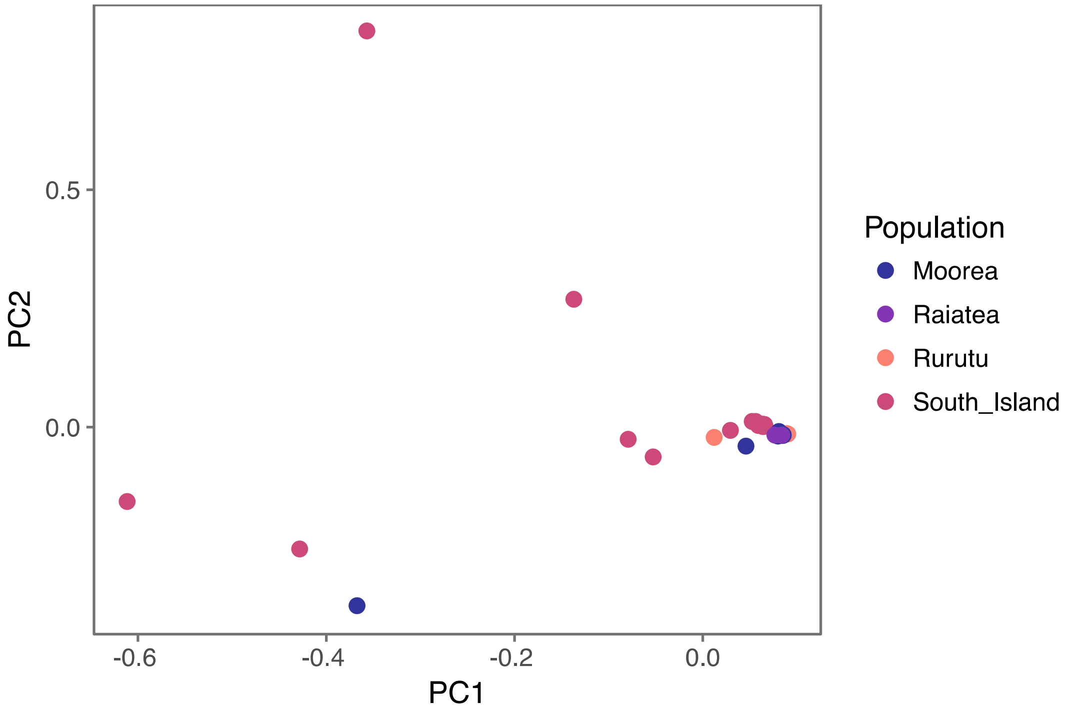


**Figure S2: Principle component analysis of genetic variation.** Based on 129,505 SNPs across 37 individuals sampled across three French Polynesian islands and South Island, New Zealand. The variance explained by PC1 and PC2 is 12.99% and 9.01%, respectively.
